# Experimental increases in temperature mean and variance alter reproductive behaviors in the dung beetle *Phanaeus vindex*

**DOI:** 10.1101/2021.02.25.432276

**Authors:** William H. Kirkpatrick, Kimberly S. Sheldon

## Abstract

Temperature profoundly impacts insect development, but plasticity of reproductive behaviors may mediate the impacts of temperature change on earlier life stages. Few studies have examined the potential for behavioral plasticity of adults to buffer developing offspring from warmer, more variable temperatures associated with climate change. We used a field manipulation to examine whether the dung beetle *Phanaeus vindex* alters breeding behaviors in response to climate change and whether adult behavioral shifts protect offspring from temperature increases. Dung beetles lay eggs inside brood balls made of dung that are buried underground for the entirety of offspring development. Depth of the brood ball impacts the temperatures offspring experience with consequences for beetle development. We placed females in either control or greenhouse treatments that simultaneously increased temperature mean and variance. We found that females produced smaller brood balls but buried them deeper in the greenhouse treatment, suggesting burial depth may come at a cost to brood ball size, which can impact offspring nutrition. Despite being buried deeper, brood balls from the greenhouse treatment experienced warmer mean temperatures but similar amplitudes of temperature fluctuation relative to the controls. Our findings suggest adult behaviors may buffer offspring from increased temperature variation due to climate change.

## 1. Introduction

Increases in temperature mean and variance associated with climate change can greatly impact the physiology and ecology of ectotherms (Angilletta, 2009). However, phenotypic plasticity, particularly behavioral plasticity, could play a key role in helping organisms cope with stressful temperatures; behavior can respond rapidly to environmental changes, potentially buffering organisms from warmer and more variable temperatures (Telemeco et al., 2009; Sih et al., 2010; Huey et al., 2012; Snell-Rood, 2013; Zuk et al., 2014; Buckley et al., 2015; Muñoz & Losos, 2018). Importantly, behavioral plasticity of adults during reproduction may protect offspring from unfavorable conditions (Refsnider & Janzen, 2012; Snell-Rood et al., 2016; Macagno et al., 2018). Understanding the capacity for reproductive plasticity is thus important for predicting how organisms will respond to climate change (Telemeco et al., 2017).

In ectotherms, reproductive behaviors of adults greatly influence the thermal environment in which offspring development takes place with consequences for phenotype and fitness (Telemeco et al., 2017; Macagno et al., 2018; Holley & Andrew, 2019; Mamantov & Sheldon, 2021). The temperatures experienced during development have profound effects on ectotherm metabolism, development rate, and body size of adults (Kingsolver et al., 2004; Ragland et al., 2008; Woods, 2009; Klok & Harrison, 2013; Pettersen et al., 2019). In many ectotherms, early life stages (e.g., eggs) are sessile and developing offspring cannot move to more favorable microclimates (Huey et al., 2012). Adjustments by adults in nesting location can alter the developmental environment of offspring with impacts on offspring survival and fitness (Snell-Rood et al., 2016; Telemeco et al., 2017; Macagno et al., 2018; Mamantov & Sheldon, 2021). However, a critical question is whether plasticity of reproductive behaviors can modify the offspring environment enough to compensate for climate change (Telemeco et al., 2009; Refsnider et al., 2013).

We studied the breeding behavior of the dung beetle *Phanaeus vindex* Macleay, 1819 to understand 1) how simultaneous increases in temperature mean and variance associated with climate change affect breeding behaviors of adults, and 2) whether behavioral plasticity of adults could compensate for climate change by buffering offspring from warmer, more variable developmental temperatures. *Phanaeus vindex* (Coleoptera: Scarabaeinae) is a medium-sized, diurnal dung beetle that ranges in open habitats and woodlands from the upper east coast to the southern USA and west to the Rocky Mountains (Edmonds, 1994). To breed, the beetles find and mate at a dung source. Females construct a tunnel below the dung source and transport dung from the soil surface to the bottom of the tunnel. The female shapes the buried dung into a brood ball and lays a single fertilized egg in it. Once the egg hatches, the developing larva eats the dung provided by the female, going through complete metamorphosis within the brood ball (Halffter & Matthews, 1966).

Maternal behavior during reproduction, including brood ball size and burial depth, shapes the environmental conditions of the offspring with consequences for survival and fitness (Hunt & Simmons, 2000, 2002, 2004; Moczek & Emlen, 2000; Snell-Rood et al., 2016; Macagno et al., 2018; Mamantov & Sheldon, 2021). Because dung from the brood ball provides the only nourishment available to developing larvae, the size of the brood ball can affect adult body size upon emergence (Lee & Peng, 1981; Moczek & Emlen, 1999; Shafiei et al., 2001; Kishi & Nishida, 2006). Importantly, smaller individuals tend to have reduced fecundity and competitive ability relative to larger individuals (Hunt & Simmons, 2000). Burial depth impacts the temperature experienced by offspring in the brood ball; offspring in brood balls near the soil surface experience warmer, more variable temperatures than those at greater soil depths (Snell-Rood et al., 2016). In ectotherms, warmer temperatures are associated with faster development rates, smaller body sizes, and reduced survival (Chown & Terblanche, 2006; Angilletta, 2009; Colinet et al., 2015; Mamantov & Sheldon, 2021). However, few studies have examined the capacity for behavioral plasticity to mediate the types of temperature changes occurring with climate change (Refsnider et al., 2013; Telemeco et al., 2017). Specifically, climate change is leading to warmer temperatures, but also more variable temperatures, which will likely be more challenging than shifts in mean temperature alone (Folguera et al., 2011; Paaijmans et al., 2013; Thompson et al., 2013; Vasseur et al., 2014; Sheldon & Dillon, 2016).

We conducted a field experiment to examine the plasticity of nesting behaviors of *P. vindex* in response to climate change by simultaneously increasing temperature mean and variance using mini-greenhouses. Our goals were to examine 1) if and how females altered reproductive behaviors, including the size, burial depth, and number of brood balls, in response to warmer and more variable temperatures, and 2) whether behavioral shifts of females buffered offspring from warmer and more variable temperatures.

## 2. Methods

### (a) Beetle collection

During two different trapping sessions in May and June 2018, we collected adult *P. vindex* near Knoxville, Tennessee, USA (36°03′25.8″N, 84°04′19.8″W) using pitfall traps baited with cow dung. We brought males (n = ~10-15) and females (n = ~20) from each trapping session back to the lab and held them in a large plastic container (18 gallon; 59 L x 46 W x 38 H cm) filled with a 4:1 mixture of topsoil:sand and covered with flexible fiberglass screen. We placed the container near a window to provide natural light, kept the room temperature at ~23° C, and fed beetles ad libitum autoclaved cow dung twice weekly. We held beetles in this colony for 3-4 weeks to allow them to mature and mate (Fincher, 1973; Blume & Alga, 1976). We checked the soil in the colonies weekly to see if females had produced brood balls. Once the colony began to form brood balls, the females were considered fertilized and ready for experimental trials.

### (b) Experimental trials

We conducted the warming experiment in an open field near Knoxville, Tennessee, USA (36°1’ N, 84°3’ W) using four, 10-day trials with start dates ranging from late June to early August 2018. We arranged 12, 7-gallon buckets (50 cm high, 28.5 cm inner diameter) in two rows with 2 m in between buckets. We drilled 5 holes into the bottom of the buckets for water drainage. We buried buckets to the brim, backfilled them with soil from the field site, and compacted the soil as much as possible. The soil in each bucket was 44 cm deep, leaving 6 cm of space between the top of the soil and top of the bucket. We placed four HOBO pendant data loggers (Model: UA-001-64, Onset, Bourne, MA) in the buckets spaced 14 cm apart (i.e., 1, 15, 29, and 43 cm below the soil surface) and recorded temperatures every 1 hour.

For each trial (four total), we randomly assigned six buckets to a heated (hereafter “greenhouse”) treatment and six to a control group. For the greenhouse treatment, we designed mini-greenhouses capable of passively and simultaneously increasing temperature mean and variance to simulate climate change. The greenhouses were made of 1/16” clear polycarbonate and shaped like a cone with a 61 cm opening at the bottom and a 7 cm opening at the top. The cone shape allowed for even heating around each bucket. Based on temperature data, the greenhouse cones simulated the types of changes expected under climate change, with an average increase of 2° C at the soil surface relative to the control buckets, though surface temperature differences could reach 4° C (Fig. 1). Temperatures in both bucket types showed natural, diurnal fluctuations and, with increasing soil depth, a decline in mean temperatures and a dampening of the amplitude of temperature fluctuation. However, temperatures at the soil depths where the data loggers were placed were warmer and more variable in the greenhouse buckets than similarly placed loggers in the control buckets (Fig. 1), thus effectively simulating conditions expected under climate change (IPCC 2013).

**Figure 1.**
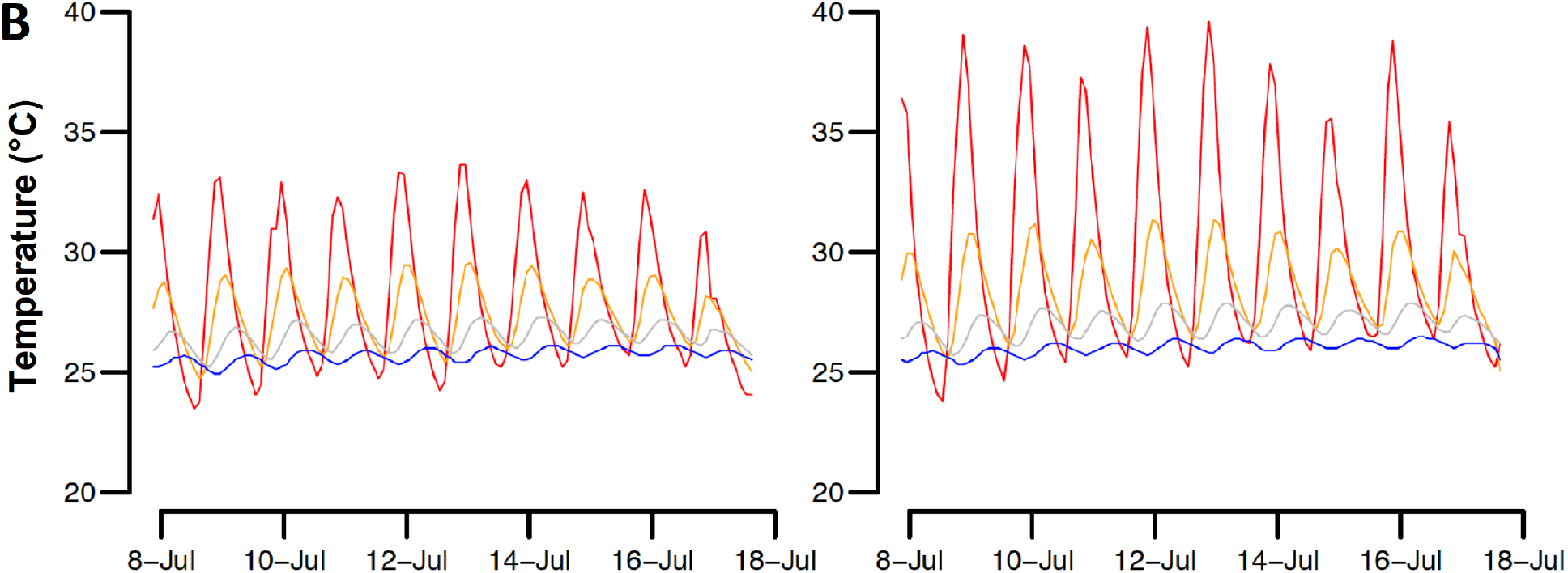
Representative temperature profiles from control (left) and greenhouse (right) buckets. Temperatures are shown from four data loggers placed at different soil depths in control and greenhouse buckets over nine days in July 2018 at the study site in Tennessee. Colors represent temperatures at different depths below the soil surface, including 1 cm (red), 15 cm (orange), 29 cm (gray), and 43 cm (blue).

For trials, we removed females from the colony, recorded mass, and randomly assigned them to a bucket. At the start of a trial, we added one female to each bucket, allowing us to eliminate the confounding effects of male behavior. Fertilized females store seminal fluid and will construct tunnels and create brood balls in the absence of males. After placing a female in a bucket, we put flexible fiberglass screen over the top secured with a bungee cord and followed by a large piece of ½ inch welded galvanized hardware cloth secured with stakes in the ground. The screen and wire prevented beetles from escaping and small animals from disturbing the trials while also allowing for more natural conditions. For warming treatments, we then placed greenhouses over the buckets. We provided beetles with ~ 25g of autoclaved cow dung at the start of the trial and then every two days thereafter to ensure they had enough for consumption and continuous brood ball construction. During these feedings, we removed all dung left on the soil surface from the previous feeding before placing new dung.

After ten days, we carefully went through the soil to uncover brood balls. We recorded the mass, burial depth, and number of brood balls from each bucket. We also removed the four temperature loggers and downloaded data. If a female did not produce brood balls, we did not use her in subsequent trials and she was excluded from statistical analyses. If a female produced brood balls, we used her in subsequent trials, typically, but not always, switching her from either a greenhouse treatment to a control or vice versa. Fifty-two percent of females produced in both bucket types. One bucket did not drain well during multiple trials, and we removed it from further analyses.

### (c) Statistical analysis

To test for behavioral shifts, we fit linear mixed effects (LME) models in R (package lme4, R version 3.6.0, R Development Core Team 2019) with the response variables of brood ball mass, burial depth, or number and the fixed effect of treatment type (greenhouse or control bucket). We included female mass as a covariate since body size may affect how much dung she can sequester (Lee & Peng, 1981). We included the random effects of trial and beetle identification to account for the non-independence of brood balls from the same trial and female, respectively (Crawley, 2007), and used model selection to decide on the random effects structure following Zuur et al. (2009). In the models for brood ball number, we removed buckets where a female did not produce a brood ball since she was not likely fertilized or where a female produced brood balls but died during the trial. Null models included an intercept and the random effects of trial and beetle identification. We selected the best-fit model considering AIC_c_ (Akaike information criterion for small sample sizes) values and the normality of residuals (Burnham & Anderson, 2002; Zuur et al., 2009; Symonds & Moussalli, 2011). Specifically, we calculated the Akaike weight of each model (wAIC_c_), which estimates the probability that the model is the best model among the candidate models considered (Burnham & Anderson, 2002). Because our model for burial depth that included the fixed effect of treatment type (i.e., control or greenhouse) was only marginally better than the null model (see Table S1), we also fit a model using the fixed effect of surface temperature recorded in each bucket rather than treatment type and all other variables described above. This allowed us to account for differences in temperatures among different buckets of the same treatment.

To examine whether behavioral shifts of adults affected the development temperatures of offspring, we used temperatures from the data loggers in each bucket to calculate temperatures experienced by each brood ball. Specifically, we fit a linear model to temperatures between neighboring pairs of data loggers to predict the mean and standard deviation of temperature for every brood ball based on its burial depth. To test whether offspring in the different treatments experienced different temperatures, we fit LME models with the response variables of either temperature mean or standard deviation at the location where the brood ball was placed and the fixed and random effects as described above.

For the final models, we computed the proportion of variance explained by fixed effects (*R*^2^_m_; Nakagawa & Schielzeth, 2013) using the r.squaredGLMM function (package MuMIn).

## 3. Results

Our goals were to examine whether females altered reproductive behaviors in warmer, more variable temperatures and whether behavioral shifts of females buffered offspring from temperature changes. Over the four 10-day trials, 21 females produced 84 brood balls. Beetles produced smaller brood balls in the greenhouse treatment (Fig. 2A); the best-fit model for brood ball size included the type of treatment as the only covariate (*β*_greenhouse_ = −13.24; wAIC_c_ = 0.61; *R*^2^_m_= 0.21; Table S1). In the second-best model that accounts for the effect of female mass, the effect of the greenhouse treatment was consistent with that of the best-fit model (*β*_greenhouse_ = −13.50; wAIC_c_ = 0.39). Beetles also buried brood balls deeper in the greenhouse treatment (Fig. 2B), although this model (*β*_greenhouse_ = 3.80, wAIC_c_ = 0.31; *R^2^_m_*= 0.05) was only marginally better than the null (wAIC_c_ = 0.28; Table S2). However, when we examined burial depth in response to the mean surface temperature of the buckets, the best-fit model that included mean surface temperature (*β*_surface temperature_ = 2.69, wAIC_c_ = 0.47; *R*^2^_m_ = 0.19) was much better than the null (wAIC_c_ = 0.13; Table S3; Fig. S1). During the 10-day trials, females produced an average of 2.5 brood balls, and we found no difference in the number of brood balls produced in the greenhouse and control buckets (Table S4; Fig. S2).

**Figure 2.**
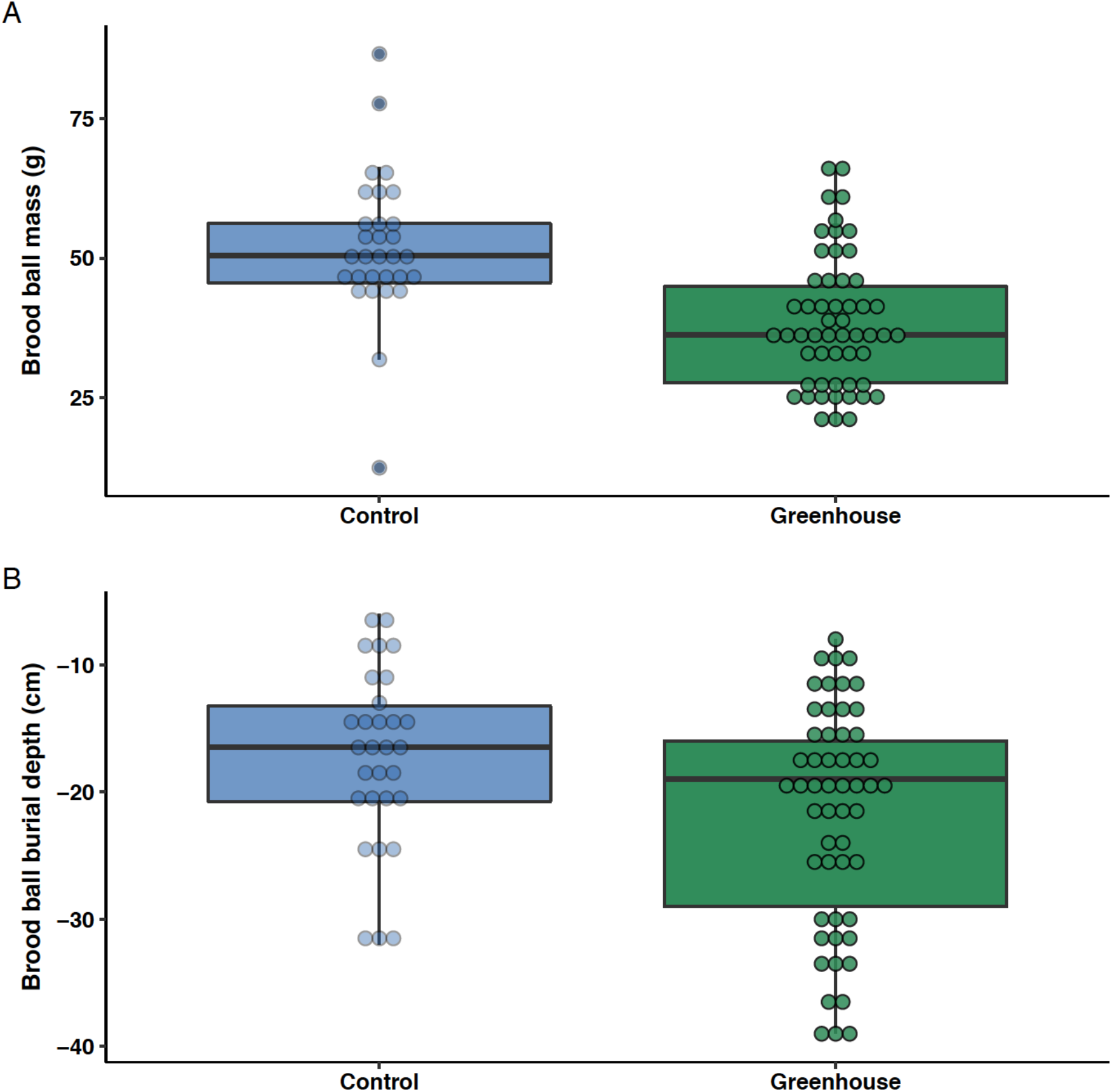
Plasticity of reproductive behaviors of female *Phanaeus vindex* dung beetles. Females in the warming treatment produced smaller brood balls (A) that were buried deeper in the soil (B) compared with beetles in the control treatment. Points show individual brood balls. Boxes show the median and first and third quartiles and whiskers show the minimum and maximum values of either brood ball size (g) or brood ball burial depth (cm).

In greenhouse buckets, the brood balls were located in areas with higher mean temperatures compared with control buckets (Fig. S3), but brood balls were located in areas with similar temperature variation regardless of treatment (Fig. S3). The best-fit model for mean temperature where brood balls were placed included the type of treatment and the effect of female mass (*β*_greenhouse_ = 0.72, wAIC_c_ = 0.54; *R*^2^_m_ = 0.17) (Table S5). In the second-best model that included just the type of treatment, the effect of the greenhouse treatment was consistent with that of the best-fit model (*β*_greenhouse_ = 0.69; wAIC_c_ = 0.46). We found no effect of treatment on temperature variation where brood balls were placed (Table S6).

## 4. Discussion

We demonstrate that female *Phanaeus vindex* altered their breeding behaviors in response to simulated climate change by producing smaller brood balls that were buried deeper in the soil. This plasticity in burial depth did not fully compensate for warming temperatures. Despite being buried deeper in the soil, brood balls from the greenhouse treatment were placed in warmer mean temperatures. Warmer temperatures during offspring development can result in faster development rates, smaller adult body sizes, and depending on the temperature, lower survival (Macagno et al., 2018; Pettersen et al., 2019; Mamantov & Sheldon, 2021). However, the mean temperatures experienced by *P. vindex* brood balls in the greenhouse treatment (26.6 °C) and control (26 °C) buckets are well below the temperatures that lead to high rates of mortality in other dung beetle offspring (Macagno et al., 2018; Mamantov & Sheldon, 2021).

In contrast to mean temperatures, *P. vindex* offspring in brood balls experienced similar amplitudes of temperature fluctuation in the greenhouse treatment and control buckets. Increased temperature variation associated with climate change may be more stressful than shifts in mean temperature alone (Vasseur et al., 2014) and could impact fitness by leading to smaller body sizes in dung beetles (Carter & Sheldon, 2020). As an example, offspring of O. taurus were three times smaller as adults when exposed to greater temperature fluctuations (Fleming et al., in press). Thus, plasticity of nest depth in *P. vindex* may buffer offspring from increased temperature variation that could otherwise negatively impact fitness (Snell-Rood et al., 2016).

Increased nest depth may protect dung beetle offspring from stressful temperatures, but time spent on tunnel construction could impact other reproductive traits (Macagno et al., 2018). *P. vindex* beetles in the greenhouse treatment produced the same number of brood balls as those in the control buckets, but the brood balls were smaller in the greenhouse treatment. One possibility for these observations is that increased nesting depth of *P. vindex* may come at a cost to brood ball size, a fitness-linked trait (Lee & Peng, 1981; Moczek & Emlen, 1999; Shafiei et al., 2001; Kishi & Nishida, 2006). Negative relationships between reproductive behaviors in response to temperature changes have been observed in other dung beetle species. *O. taurus* beetles exposed to warmer mean temperatures (Mamantov & Sheldon, 2021) and increased diurnal temperature variation (Holley & Andrew, 2020) did not alter brood ball size, but they produced fewer brood balls. Interestingly, *O. hecate* appeared to show no tradeoff in reproductive behaviors in response to warming; individuals produced more brood balls of the same size and buried them deeper in response to warmer mean temperatures (Mamantov & Sheldon, 2021). However, fitness could be impacted in other ways, such as via reduced egg production and quality. As an example, in the ball-rolling dung beetle *Sisyphus rubrus*, warmer treatments did not affect the size or number of brood balls produced, but beetles buried fewer brood balls, which are more likely to contain eggs than unburied brood balls (Holley & Andrew, 2019). Thus, shifts in one behavior in response to temperature changes may come at a cost to other fitness-linked traits.

Is the behavioral plasticity we observed in *P. vindex* adaptive? Though we did not examine fitness, we did find potential tradeoffs that suggest an adaptive response. Specifically, brood balls were smaller in the greenhouse treatment, which can reduce fitness via impacts on adult body size (Lee & Peng, 1981; Moczek & Emlen, 1999; Shafiei et al., 2001; Kishi & Nishida, 2006). However, by digging deeper in the soil, beetles were able to buffer their offspring from potentially stressful temperatures. This suggests that behavioral plasticity to temperature change may be adaptive in *P. vindex* by increasing offspring survival even if the reduced brood ball sizes result in smaller individuals. Though behavioral plasticity may be adaptive in *P. vindex*, this is not the case for all ectotherms. For example, *Sceloporus tristichus* lizards reduced nesting depth in response to warmer temperatures, thus increasing heat stress on embryos (Telemeco et al., 2017). Understanding the behavioral adjustments made by ectotherms in response to temperature change is a key step to predicting the potential impacts of climate change (Huey & Tewksbury, 2009).

## Acknowledgements

We are grateful to Susan Riechert and Mac Post for providing access to their property for the field experiment. We thank Matt McGee and Morgan Fleming for field assistance, and Luis Carrasco and Xingli Giam for helpful comments on an earlier draft.

## Data accessibility

Data are accessible as electronic supplementary material.

## Authors’ contributions

KSS conceived of the study and designed the experiment. WHK performed the experiment. KSS analyzed data with assistance from WHK. Both authors wrote the manuscript, gave final approval for publication, and agree to be held accountable for the work performed therein.

